# A novel biomechanical model of the mouse forelimb predicts muscle activity in optimal control simulations of reaching movements

**DOI:** 10.1101/2024.09.05.611289

**Authors:** Jesse I. Gilmer, Susan K. Coltman, Geraldine Cuenu, John R. Hutchinson, Daniel Huber, Abigail L. Person, Mazen Al Borno

## Abstract

Mice are key model organisms in neuroscience and motor systems physiology. Fine motor control tasks performed by mice have become widely used in assaying neural and biophysical motor system mechanisms. Although fine motor tasks provide useful insights into behaviors which require complex multi-joint motor control, there is no previously developed physiological biomechanical model of the adult mouse forelimb available for estimating kinematics nor muscle activity or kinetics during behaviors. Here, we developed a musculoskeletal model based on high-resolution imaging of the mouse forelimb that includes muscles spanning the neck, trunk, shoulder, and limbs. Physics-based optimal control simulations of the forelimb model were used to estimate *in vivo* muscle activity present when constrained to the tracked kinematics during reaching movements. The activity of a subset of muscles was recorded and used to assess the accuracy of the muscle patterning in simulation. We found that the synthesized muscle patterning in the forelimb model had a strong resemblance to empirical muscle patterning, suggesting that our model has utility in providing a realistic set of estimated muscle excitations over time when given a kinematic template. The strength of the similarity between empirical muscle activity and optimal control predictions increases as mice performance improves throughout learning of the reaching task. Our computational tools are available as open-source in the OpenSim physics and modeling platform. Our model can enhance research into limb control across broad research topics and can inform analyses of motor learning, muscle synergies, neural patterning, and behavioral research that would otherwise be inaccessible.

**NEW & NOTEWORTHY:** Investigations into motor planning and execution lack an accurate and complete model of the forelimb, which could bolster or expand on findings. We sought to construct such a model using high detail scans of murine anatomy and prior research into muscle physiology. We then used the model to predict muscle excitations in a set of reaching movements and found that it provided accurate estimations and provided insight into an optimal-control framework of motor learning.

## INTRODUCTION

*Mus musculus* (Mice) are key model organisms for behavioral studies in neuroscience and physiology, including for tasks that assay fine motor control and motor learning. Mice can perform tasks such as manipulandum control (Bollu et al., 2019), dexterous reach (Becker et al., 2020, Fleischer et al., 2023; Khanna et al., 2021; Yang et al., Wagner et al., 2021), and can learn complex behaviors with and without training (Burgess et al. 2017, Serradj et al., 2023, Sauerbrei et al., 2020, Galinanes et al., 2018, Conner at al., 2021). Mice are a useful organism for studying human disease, and their behaviors reflect an evolutionarily preserved trait applicable to human motor disorders (Iwaniuk 2000). They also convey a benefit to researchers because of their accessible genetics and a slate of powerful molecular tools, which can allow researchers to perturb neural behavior through optogenetics (Deisseroth et al., 2006).

However, despite the utility of mice as a model organism in motor learning, there are no high-resolution reconstructions of the adult mouse forelimb, nor are there any physiological biomechanical models of the mouse forelimb that incorporate fully developed muscle morphology. Biomechanical models are useful tools for motor systems and neuromechanics researchers that can provide detailed insights into muscle activity and limb kinematics (e.g., fiber length and velocities) that would otherwise be difficult or impossible to access through empirical observations. State-of-the-art recording methods can only measure the activity of 3-4 muscles of the 25+ muscles in the murine forelimb (Zia et al., 2020; Chung et al., 2023). Therefore, we set out to construct and evaluate a model of the mouse forelimb, which would be a valuable tool for researchers studying dexterous behaviors in mice.

The only currently available mouse forelimb model, developed recently in a full-body mouse model (Ramalingasetty et al., 2021), was based on mouse embryo data (Delaurier et al., 2008), and lacks many of the large muscles originating from the scapula. Ramalingasetty and colleagues reported that modeling the mouse forelimb was challenging and identified improving their forelimb model as a remaining challenge for future work. Moreover, reference books on limb anatomy present two-dimensional (2D) illustrations of the limb musculature (Hebel 1986), which makes it challenging to extract accurate locations of the attachment points and the three-dimensional (3D) tissue paths from these references (Delaurier et al., 2008). Attachment points have been shown to be the most important factor in estimating how effective a muscle is in producing a joint rotation or moment (Charles et al., 2016). Computing these quantities directly from dissections is challenging because of the size of the forelimb muscles, whose tendon insertion points are separated by as little as tens of microns.

We were able to more accurately identify the muscle attachment sites and muscle paths than previously possible using mouse and rat atlas data through the use of light sheet microscopy. Building on the detailed description of muscle anatomy integrated into a hindlimb biomechanical model by Charles et al. (2016), we aimed to extend this work by developing a similar model for the forelimb using imaging data. We started by scanning and recreating the forelimbs of two adult mice. Muscles with insertions onto the humerus originate from sites that span most of the mouse’s trunk and spine, necessitating imaging of much of the mouse body. We limited our reconstruction to muscles that had insertions onto the humerus, radius, and ulna, as reconstruction of muscles with insertions onto the scapula and those that inserted onto the hand was infeasible given the resolution of imaging performed. Once the muscles had been traced and reconstructed, they were used to construct the musculoskeletal geometry of the biomechanical model, including attachments and lines of action, using the OpenSim modeling and physics simulation environment (Delp et al., 2007; Kewley et al., 2024). We used published results on mouse forelimb muscle architecture to set the muscle parameters (Mathewson et al., 2012). With a model based on highly accurate reconstructions and mouse physiology, we then hypothesized that our model would be well suited to the prediction of muscle activity during motor behaviors such as skilled reaching.

To evaluate the utility of the model, we then aimed to replicate physiological kinematics and predict simultaneously recorded muscle activity. We analyzed a subset of a dataset comprising thousands of reaches from three mice, which included both 3D kinematics and the activity of a subset of muscles involved in the reaching movements (biceps, triceps long head and triceps lateral head electromyography (EMG)). The empirical kinematics were used as constraints when producing muscle-driven synthesized kinematics with optimal control in OpenSim. The empirical EMG was then used as a ground truth for comparison against the optimal control EMG predictions produced in simulation.

We found that optimal control-based simulations using the model were able to recreate reach kinematics accurately using synthesized muscle excitations. When examined against empirical EMG, synthetic EMG closely resembled the mean activation during kinematically matched reaches. These results suggest that our model is capable of replicating realistic reach kinematics and muscle activity. Additionally, we had access to behavioral data that spanned the extent of learning, thus we performed analyses that reveal that the optimal control solutions are closer to the empirical solutions (i.e., the patterns employed by real mice) as reaching performance improves throughout learning. In other words, mice may employ muscle patterning solutions that more closely resemble optimal control solutions as they become more skilled at the task.

More broadly, this model should provide insight into forelimb behaviors that would otherwise be inaccessible by experimental means, and we hope that access to a robust description of the forelimb’s kinematics, forces, and muscle activity will advance understanding of mouse behavior. Our computational tools are available as open-source for researchers interested in analyzing muscle activity during mouse forelimb movements.

## METHODS

### Animals

All procedures followed National Institutes of Health Guidelines and were approved by the Institutional Animal Care and Use Committee at the University of Colorado Anschutz Medical Campus under protocol #43. Animals were housed in an environmentally controlled room, kept on a 12-h light– dark cycle and had ad libitum access to food and water, except during behavioral training and testing as described below. Adult C57BL/6 (Charles River Laboratories) mice of either sex (3 females and 1 male) were used in behavioral experiments. The animals for the light sheet imaging were part of experiments approved by the Animal Care Committee of the University of Geneva and by the veterinary office of the “Direction générale de la santé” of the Canton of Geneva. Adult C57BL/6 mice of the female sex (2 Females) were used in imaging experiments.

### Anatomical high-resolution imaging

Accurate prediction of muscle activity during movements is predicated on a sufficient description of the underlying anatomy and physiology. With the goal of creating a predictive model, we began by collecting anatomical data. We obtained 3D scans of mouse forelimbs and trunk musculature through large scale light sheet microscopy imaging of two wild type female mice (11 weeks old). The dataset contains imaging that captured the left distal shoulder and proximal forelimb (Mouse A), the right distal forelimb and paw (Mouse A), and both forelimbs, shoulders, and trunk (Mouse B). Only the left shoulder, trunk, and proximal forelimb were reconstructed in Mouse B. Mouse A and B are different from the four mice used in the empirical reaching dataset with EMG. This imaging dataset provides detailed anatomical information to inform our model development.

### Mouse and tissue preparation

Mice were euthanized via subcutaneous injection of pentobarbital. Transcardiac perfusion with saline, followed by 4% paraformaldehyde (PFA) and 0.01% heparin, was performed to preserve tissue integrity. The circulatory system was then washed with saline solution, and the skin was removed. To clear the entire mice, a chamber was set up in the manner that the whole bodies were immersed under the solution which was perfused in a closed loop through the vascular system using a peristaltic pump. All the following incubation steps were performed through this perfusion set up. The bone structures of the mice were decalcified by incubating them in 20% EDTA at 37°C for ten days, with the EDTA solution being renewed every 3 days.

Imaging subjects were pretreated with methanol following the principle of iDISCO+ tissue clearing protocol <u>(Branch et al., 2019</u>; Renier et al., 2014). The tissue was introduced to a gradually increasing concentration of methanol, starting with 20% and increasing by 20% every hour. The clearing chamber was maintained at room temperature. After 5 hours of exposure to methanol, the tissue was chilled at 4° C overnight and then bathed in 66% dichloromethane (DCM) and 33% methanol for 24 hours. The tissue was then bathed twice in 100% methanol for 1 hour before being chilled for 1 hour at 4°C and then transferred into 5% hydrogen peroxide in methanol for 48 hours. Finally, tissue was rehydrated through 1 hour immersion in 80%/60%/40%/20% methanol for one hour per 20% decrement, then transferred to 1x PBS for 24 hours, followed by immersion in a 100ml PBS 10x and 2 ml TritonX-100 solution that was filled to 1L with distillate water. Since only the tissue autofluorescence was targeted for imaging, no antibodies were used in this clearing step process. The tissue was permeabilized with a solution (500 mL) consisting of 400 mL PTx.2, 11.5 g glycine, and 100 mL dimethylsulfoxide (DMSO). The tissue was bathed in solution for 4 days, then transferred to a blocking solution of 42 mL PTx.2, 3 mL donkey serum, and 5 mL DMSO for 3 days. Finally, tissue was washed with 100 mL PBS 10X, 2 mL Tween-20, 1 mL of 10mg/mL heparin, and filled to 1L with distillate water. The tissue was then redehydratedthrough preparation in 20%/40%/60%/80%/100% methanol in 1-hour steps, then bathed in 100% methanol overnight. Afterwards, the tissue was bathed in 66% DCM and 33% methanol for 4 hours, then in 100% DCM for 15 minutes twice, in succession, and finally immersed in DBE solution for imaging.

### Imaging parameters

Dissected mouse limbs were arranged in a prone position prior to imaging. Scans were taken with 8.23 um per pixel scans at 8x zoom, with 5 um steps in the z-plane. Imaging was performed using mesoSPIM (Voigt, et al., 2019). Immunostaining was captured in the green channel (488 nm laser) and was imaged using mode tiling wizard with an offset by 75% and a filter set to 530/43. Mouse A’s forearms were dissected and imaged in their entirety. Mouse B was imaged from the base of the skull through the joint of the femur and tibia and the entirety of the depth of the sample.

### Anatomical segmentation and reconstruction

Muscle segments were manually reconstructed in 3D. We found that muscle density and striation was a sufficient marker of muscles to identify them with light sheet microscopy using the tissue autofluorescence in the green channel, which were enhanced through immunostaining without antibodies (see **Methods: Mouse and tissue preparation**). We used the raw imaging of the mouse anatomy and segmented individual muscles into 3D shape objects using 3D Slicer (Federov et al., 2012) (Fig. 1A). We also segmented the forelimb bones to obtain landmarks and geometries for use in the model. Because not every data set had complete data for the entire forelimb, right and left anatomical datasets were combined through manual alignment in Blender (Blender D.T., 2022). We used anatomical landmarks on the humerus, ulna, and radius to align muscle reconstructions, as these bones were present in all three imagining datasets (left and right forelimbs of mouse A, and whole body of mouse B using the left forelimb). The reconstructions were scaled according to the radius of the bones and confirmed visually by examining the degree of overlap between reconstructions.

**Figure 1.**
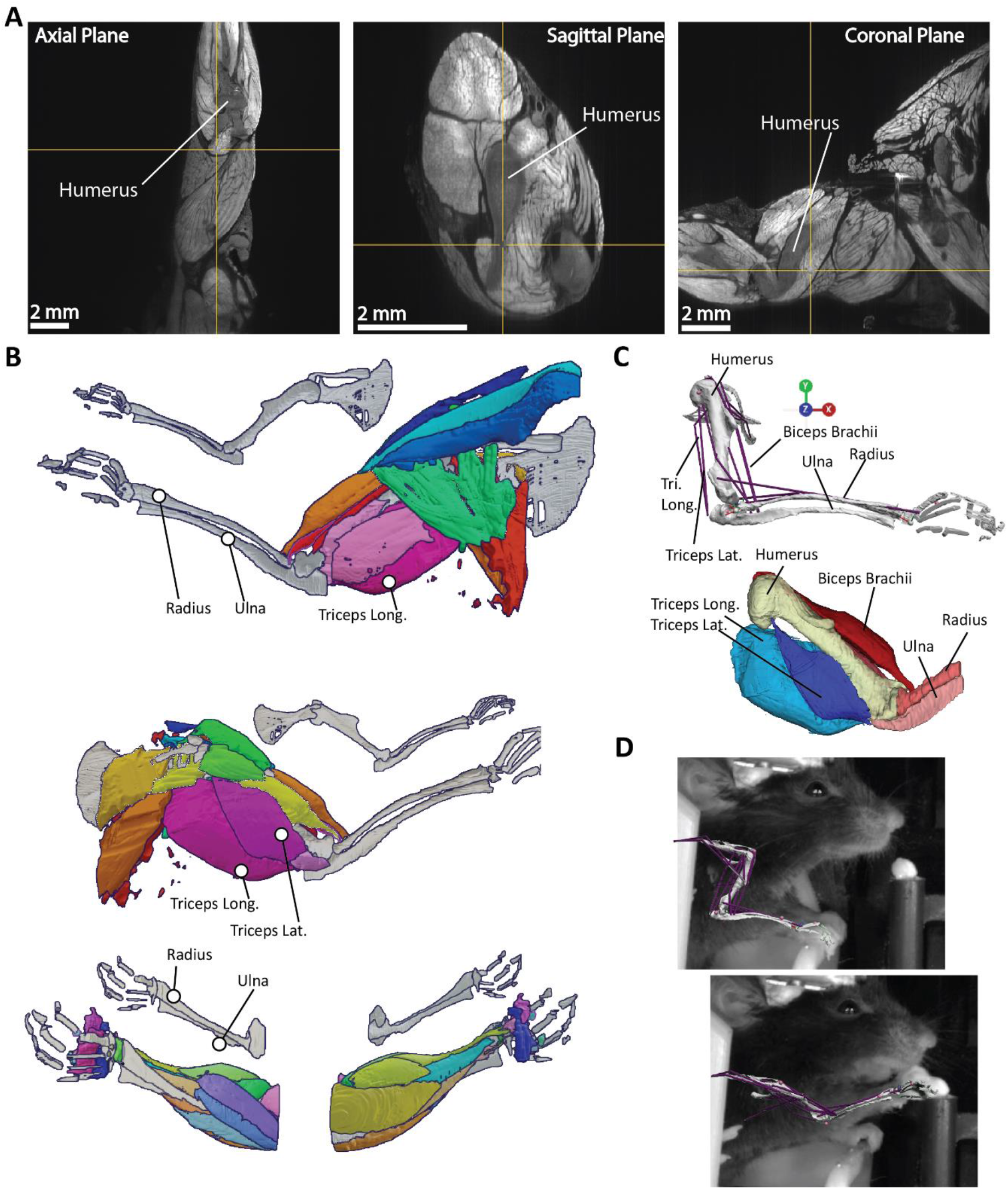
Anatomical reconstruction. **A**. Optical slices of the mouse forelimb in the axial, sagittal, and coronal planes. The mouse arm is oriented in the prone position. Labels added to highlight prominent muscles as an example of a reconstruction target. **B**. 3D projections of optical tracing results as a composite across mice. Upper panels show composite scan, while lower panels show the left hand of mouse A to highlight density of wrist-inserting muscles. 3D projections show morphology and attachment sites of muscles on bones that were used to create biophysical model. **C**. Biomechanical model (OpenSim) reconstruction developed from the 3D projections. **D**. Biomechanical model projection on video images taken from of a mouse reaching.

### Development of a biomechanical model of the mouse forelimb

With a detailed set of reconstructions, we next sought to leverage anatomical descriptions to construct a biomechanical model in OpenSim, a widely-used physics-based modeling and simulation environment used to study movements of humans and other species. The anatomical model was assembled using OpenSim Creator (Kewley et al., 2024). We have also converted the model in MuJoCo (Todorov et al., 2012), but that model was not used for simulations in this study. Each individual muscle’s parameters were derived from optical measurements and from previous parameters in Mathewson et al., 2012 and Charles et al., 2016. These were used to set the biophysical properties of the modelled muscles. We used De Groote-Fregly (De Groote et al., 2016) Hill-type muscles within the model, and opted to ignore complex tendon dynamics (i.e., using rigid tendons with no force-length/velocity properties), both to facilitate the production of a functional model and because we did not have access to sufficient data regarding tendon physiology purely from imaging data. Our Hill-type muscles were parametrized by four parameters (maximum isometric force, optimal fiber length, tendon slack length and pennation angle). All these parameters, except for tendon slack length, were determined experimentally in the muscle dissection study of Mathewson et al., 2012 and through interpolation from known values when a muscle was not described in prior literature. The tendon slack length parameter represents the length where a tendon develops passive elastic force (Uchida & Delp 2021). This parameter cannot be measured experimentally and was set using the optimization procedure of Buchanan et al., 2004, as is commonly performed in the field (Charles et al., 2016), assuming that muscle fibers remain within 0.5 to 1.5 times optimal fiber length throughout the joint’s range of motion, which were estimated from both anatomical constraints and video of mouse behavior. Based on the muscle paths from the digital segmentation, we used wrapping surfaces, which are geometric objects in OpenSim, to constrain the muscles to have realistic paths of action. This is necessary for the model to produce realistic moment arms (Charles et al., 2016). We set other parameters in the muscles such as the maximum contraction velocity, the activation time constants and the force-length curves scaled based on prior work on mouse physiology (Charles et al., 2016; see our open-source model for details). We calculated the physiological cross-sectional area (PCSA) by the standard formula developed by Alexander & Vernon (1975), that is, muscle volume divided by fiber length. Muscle fiber pennation angle is set separately in OpenSim models; thus not directly used in PCSA calculations. Bone volume was determined in reconstruction and was uniformly multiplied by a murine bone density scalar (.00425 kg/cm^2^), determined from a literature search for empirical measures (Robbins et al., 2018) and prior models of the mouse (Charles et al., 2016), as well as estimations of the center-of-mass and inertia. The resulting model has 21 muscles and 5 bones (along with a composite hand body segment), with the scapula and clavicle serving as fixed position bodies. The model includes four degrees-of-freedom: shoulder elevation, extension, and rotation, as well as elbow flexion. The model is also capable of wrist flexion and rotation, but these degrees-of-freedom were fixed during our simulations. A description of the model geometry is available in Table 1 and the muscle parameterization in Table 2.

**Table 1.** Muscle origins and insertions.

| Muscle Name | Origin Parent-Body | Origin Coordinates (m) | Insertion Parent-Body | Insertion Coordinates (m) |
| --- | --- | --- | --- | --- |
| Anconeus | Humerus | [5.2e-3, 5e-4, 1.07e-2] | Ulna | [2.4e-3, -2e-4, 8.8e-3] |
| Anconeus, Short Head | Humerus | [3.6e-3, -9e-4, 9.1e-3] | Ulna | [3.8e-3, -6e-4, 8.3e-3] |
| Biceps, Long Head | Scapula | [8.5-3, 2.3e-2, 1.2e-2] | Ulna | [2.1e-3, 4.9e-5, 8.8e-3] |
| Biceps, Short Head | Humerus | [9.1e-3, 1.5e-3, 1.2e-2] | Ulna | [2.1e-3, 9.8e-5, 8.8e-3] |
| Brachialis, Proximal Head | Humerus | [2.3e-3, 5.9e-5, 8.8e-3] | Ulna | [2.2-3, 5.9e-5, 8.8e-3] |
| Brachialis, Distal Head | Humerus | [8.2e-3, 1.4e-3, 1.2e-2] | Ulna | [2.2-3, 5.9e-5, 8.8e-3] |
| Brachioradialis | Humerus | [4e-3, 6.5e-4, 9.1e-3] | Radius | [-3e-3, 1.3e-3, 1.2e-2] |
| Deltoid, Medial | Clavicle | [9.3e-3, 5e-4, 1.3e-2] | Humerus | [5.5e-3, 1.2e-3, 1.2e-2] |
| Deltoid, Posterior | Scapula | [8.7e-3, 1.2e-3, 1.2e-2] | Humerus | [5.9e-3, 1.5e-3, 1.1e-2] |
| Flexorcarpiradialis | Humerus | [3.4e-3, -9.3e-4, 9.2e-3] | Hand | [-4e-3, 1.6e-3, 1.2e-2] |
| Infraspinatus | Scapula | [1.1e-2, 1.3e-3, 1.2e-2] | Humerus | [8.3e-3, 1.6e-3, 1.2e-2] |
| Latissimus Dorsi, Caudal | Spine* | [1.8e-2, -3e-3, 1e-2] | Humerus | [5.5e-3, 1.7e-3, 1.1e-2] |
| Latissimus Dorsi, Rostral | Spine* | [1.5e-2, -7.1e-4, 1.3e-2] | Humerus | [5.9e-3, 1.5e-3, 1.1e-2] |
| Pectoralis Major, Anterior | Rib-cage* | [9.1e-3, -1.6e-3, 1.7e-2] | Humerus | [5.3e-3, 1.4e-3, 1.2e-2] |
| Pectoralis Major, Posterior | Rib-cage* | [1e-2, -3.1e-3, 1.5e-2] | Humerus | [5.5e-3, 1e-3, 1.2e-2] |
| Pectoralis Minor, Clavicular | Clavicle | [1.1e-2, -9.6e-4, 1.3e-2] | Humerus | [5.6e-3, 1.2e-3, 1.2e-2] |
| Pronator Teres | Humerus | [3.6e-3, -6.7e-4, 9.3e-3] | Radius | [8.8e-4, 1.2e-3, 1e-2] |
| Subscapularis | Scapula | [1.3e-2, 4e-4, 1.3e-2] | Humerus | [8.2e-3, 6.2e-4, 1.2e-2] |
| Triceps, Long Head | Scapula | [9.8e-3, 1.2e-3, 1.2e-2] | Ulna | [4.2e-3, -5.7e-4, 8e-3] |
| Triceps, Lateral Head | Humerus | [8.4e-3, 1.6e-3, 1.2e-2] | Ulna | [4.1-3, -4.5e-4, 8.3e-3] |
| Triceps, Medial Head | Humerus | [6.5e-3, 2.5e-4, 1.1e-2] | Ulna | [3.8e-3, -2.3e-4, 8.4e-3] |

\* Spinal and rib attachments made to Scapula fixed ground object.

**Table 2.** Muscle Parameters.

| Muscle Name | Max Isometric Force (N) | Optimal Fiber Length (m) | Pennation Angle (rad) | Tendon Slack Length (m) |
| --- | --- | --- | --- | --- |
| Anconeus | 0.023 | 0.003 | 0.1 | 0.0002 |
| Anconeus, Short Head | 0.02 | 0.0015 | 0.1 | 0.00015 |
| Biceps, Long Head | 0.093 | 0.0085 | 0.1 | 0.0002 |
| Biceps, Short Head | 0.018 | 0.005 | 0.1 | 0.0005 |
| Brachialis, Proximal Head | 0.066 | 0.007 | 0.1 | 0.0001 |
| Brachialis, Distal Head | 0.067 | 0.007 | 0.1 | 0.0001 |
| Brachioradialis | 0.02 | 0.007 | 0.1 | 0.0001 |
| Deltoid, Medial | 0.069 | 0.006 | 0.2 | 0.0002 |
| Deltoid, Posterior | 0.068 | 0.0035 | 0.2 | 0.0001 |
| Flexorcarpiradialis | 0.02 | 0.007 | 0.1 | 0.0001 |
| Infraspinatus | 0.065 | 0.003 | 0.2 | 0.0001 |
| Latissimus Dorsi, Caudal | 0.133 | 0.011 | 0.36 | 0.0005 |
| Latissimus Dorsi, Rostral | 0.1133 | 0.011 | 0.36 | 0.0005 |
| Pectoralis Major, Anterior | 0.233 | 0.008 | 0.3 | 0.0002 |
| Pectoralis Major, Posterior | 0.170 | 0.007 | 0.3 | 0.0005 |
| Pectoralis Minor, Clavicular | 0.033 | 0.0056 | 0.25 | 0.0002 |
| Pronator Teres | 0.02 | 0.003 | 0.1 | 0.0001 |
| Subscapularis | 0.34 | 0.005 | 0.2 | 0.0001 |
| Triceps, Long Head | 0.612 | 0.008 | 0.3 | 0.0007 |
| Triceps, Lateral Head | 0.125 | 0.007 | 0.17 | 0.0001 |
| Triceps, Medial Head | 0.16 | 0.004 | 0.2 | 0.0001 |

### Model scaling

Individual mice have variable limb dimensions that models must be altered to accommodate. We accomplished this by using the scale tool in OpenSim to automatically scale the mass, length, and muscle parameters of the model to fit the observed kinematic data originating from a particular mouse subject. We used DeepLabCut (Mathis et al., 2018) to estimate paw, elbow and shoulder markers from video. Our scripts adjusted the marker positions based on a 3D skeletal model with estimated limb lengths (derived from mean inter-marker distances). These adjusted marker positions were then used to scale the OpenSim model to the mouse’s proportions.

### Mouse behavior

Mouse reaching behavior was recorded from two cameras, spaced 55 degrees apart, and then processed using the DeepLabCut 3D motion tracking software (Mathis et al., 2018). Post-processing of the data parsed the recorded movements into reaching movements. We collected data in a forelimb reaching task and recorded EMG from the biceps brachii, triceps long head, and triceps lateral head in four mice. The activity of three muscles were measured simultaneously with Myomatrix arrays (Zia et al., 2020, Chung et al., 2023), using a bipolar headstage. We estimated the elbow joint angle from the 3D markers. We used the average limb lengths to adjust the DeepLabCut paw, elbow and shoulder markers and ensure that the limb lengths remain constant throughout the video, which is necessary for accurate tracking by the model. Because our forelimb model only has rotational degrees-of-freedom on the shoulder, we could not capture the small translational movement occurring at the shoulder during head-fixed reaching. We subtracted the shoulder markers displacements from the elbow and paw markers to keep shoulder positional coordinates fixed in our simulations.

The mice underwent two surgeries, 5-7 days apart. First, a head plate was installed to enable head fixation during the experimental training and testing, followed by a second surgery to implant the EMG arrays. Mice were placed on the experimental rig 5-7 days post-EMG implant. Post-surgery, mice were maintained at 80-85% of their initial body weight throughout training and testing phases. For data collection, mice were head-fixed on the experimental rig and trained to perform a single pellet retrieval task (Figs. 1D, 2A). Kinematic measurements and EMG recordings were continuously collected throughout the entire training process, starting from the first session when the mice were completely naive, until they gradually improved and achieved expertise in the task. This training period spanned 11 to 26 sessions, with the mice making their first successful reach (i.e., retrieving a pellet) between 2 to 5 sessions on the rig.

Processed EMG envelopes were normalized to the maximum contraction recording during the session. EMG is usually normalized to the maximum voluntary contractions in studies with human subjects (Kendall et al., 2005). We rectified the EMG signals and then filtered them with a bandpass and lowpass filter suite. We bandpassed the signal from 5 to 500 Hz, rectified the signal, then low-passed further with a cutoff of 10 Hz. Additionally, we normalized the filtered EMG signals with a z-score measure. Each muscle was recorded through 4 leads, but only the qualitatively determined cleanest lead per muscle was used for this study.

### Selection of reaches for simulation

Our dataset spanned the entirety of reach training for four mice, and because of the progression of learning, there was natural variance in kinematics performed. We opted to select only from ‘expert’ mice and to use baseline EMG datasets that were derived from similar reaching kinematics, which eliminated one of the mice. We grouped reaches using the 2-norm metric on 3D paw kinematics to assess similarity, and then selected two sets of ten reaches per mice (n=3), with each set having a different kinematic profile (see Fig. 2C). We enforced expertise by selecting reaches that occurred only after the initial 4 sessions of learning, which was a typical epoch for mice to reach moderate success in reaching. We also compared the optimal control predictions between the early and late sessions of learning. Early sessions were selected from the four mice discussed above (i.e., including the mouse that did not achieve expertise status). Ten reaches were selected from each mouse for the early dataset. Early sessions were restricted to the first 3 sessions of learned reaching.

**Figure 2.**
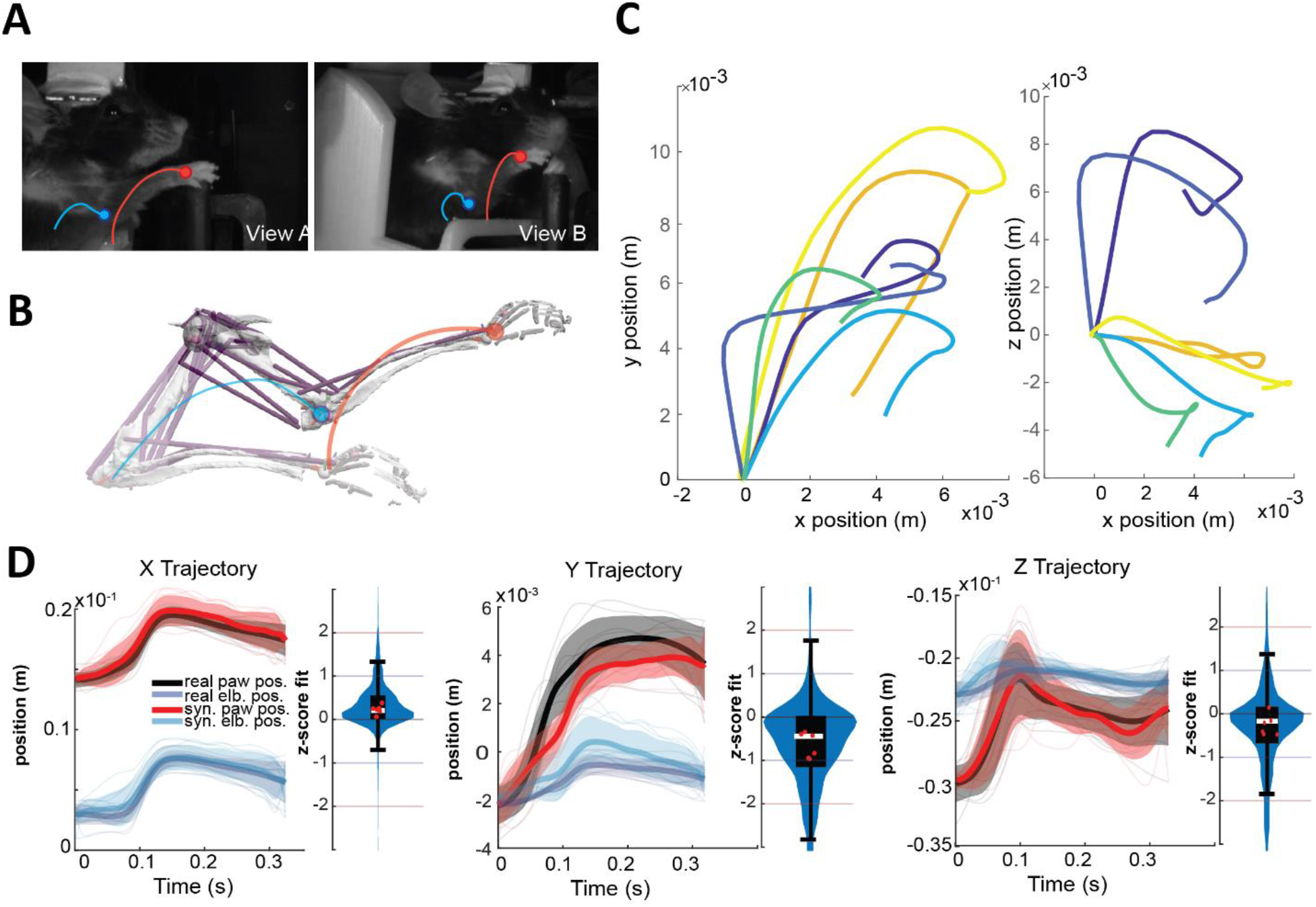
**A**. Example video screenshots with schematic elbow and paw marker trajectories. **B**. A biomechanical model with virtual markers on the elbow and paw. An optimal control problem is solved to minimize the difference between the virtual and empirical markers. **C**. Means of six sets of reaches selected for simulation, with the x vs y (forward vs vertical) dimensions plotted on the left and x vs z (forward vs lateral) on the right. **D**. Mean and standard deviation of 60 reaches for the empirical and synthesized marker trajectories. The z-score of the synthesized markers are largely within 1 standard deviation (see violin plots in blue), and the means per set of ten reaches are all within 1 standard deviation (red dots). Black box plots denote median (white bar), 25 to 75^th^ percentile distributions (black box), and 10^th^ to 90^th^ percentile distributions (short horizontal black lines).

### Optimal control

Optimization of synthetic movements matching empirical kinematics was conducted with direct collocation in Moco (Dembia et al., 2020) as it is well-suited for simulations that track experimental data (“inverse simulations”; e.g., Bishop et al., 2021). Direct collocation enforces the equations-of-motion and physiological relationships as constraints in a nonlinear optimization problem which solves for the states of the musculoskeletal system and the muscle activity over the duration of the simulation. The optimization’s objective is to minimize a cost function of two terms: one term that is a proxy for effort (i.e., the sum of muscle activations squared) and one term that represents the tracking cost (i.e., the deviation between the synthesized and the experimental kinematics). The cost function equation is:

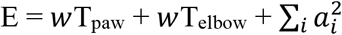

Here, T_paw_ is the 2-norm squared difference between model and experimental 3D paw coordinates. The experimental coordinates comprised of 100 timepoints during the ballistic phase of reaching. The same holds for T_elbow_, which is derived from tracking of the mouse elbow position. Term *a*_*i*_ denotes the activation of muscle *i* in the model and *w* is a scalar weight set to 10^9^. We optimized over 2500 iterations and 100 mesh points. The simulation was also constrained to start with the joint angles derived during the scaling of the model. The optimization would end early if a convergence tolerance of 1*e*^−7^ was reached. The optimization typically ran for ten minutes on a computer with specifications listed in supplemental Table 1. Rarely, MoCo would not resolve an appropriate kinematic solution for muscle driven simulations (in 2 trials), in which case the cost function was altered to *w* = 10^12^ then re-simulated.

We compared range normalized predicted muscle activity and EMG measurements using the mean absolute error (MAE) metric at an optimal lag (in a range of –50 to +10 ms; we used a lag of 0 ms for Figure 3 and the late reach set in Figure 4; early reaches had an optimal lag of –50 ms in Figure 4).

**Figure 3.**
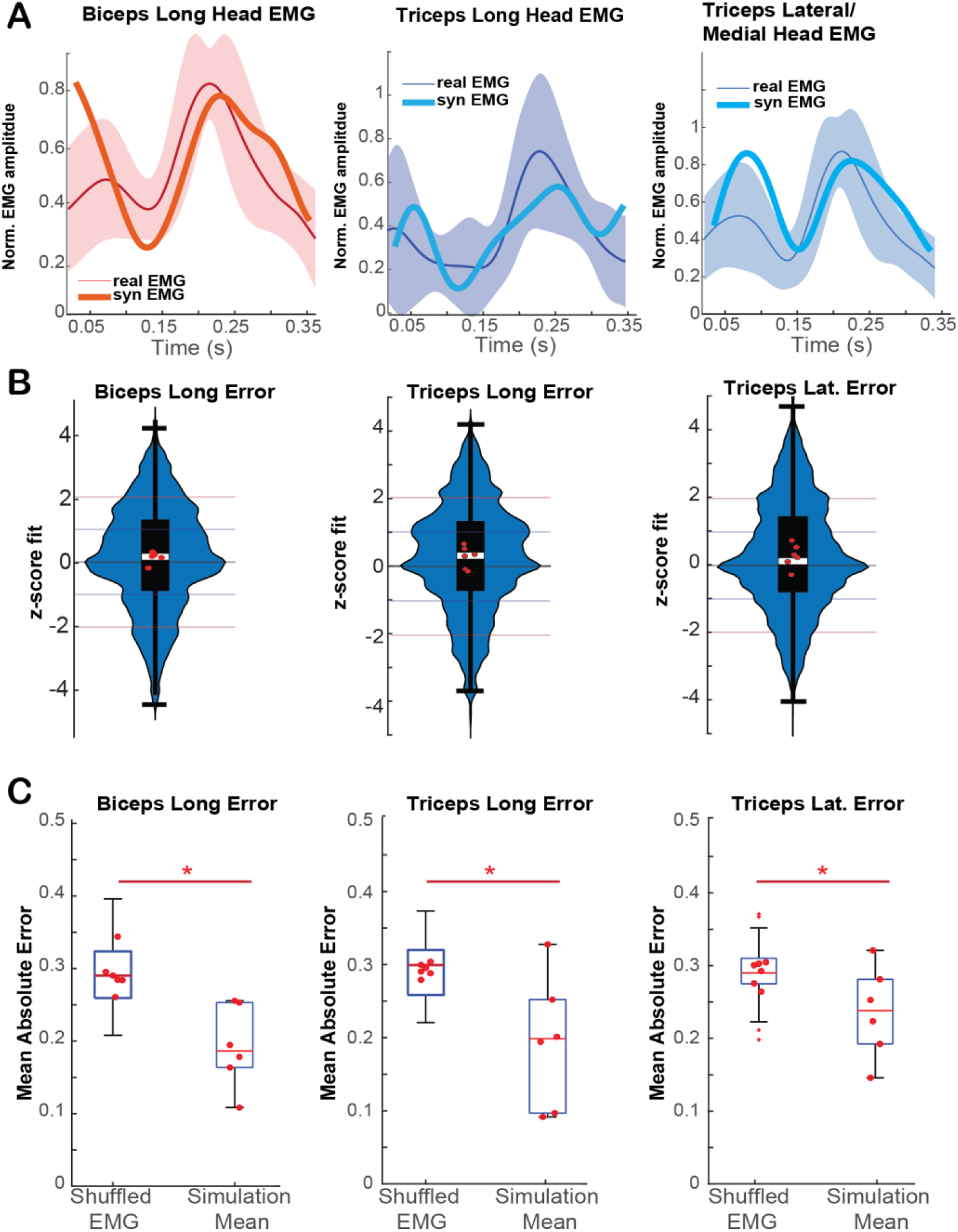
A. Comparisons of synthesized muscle excitations and experimental EMG activity. Curves show means and standard deviations (line and shaded region) of ten reaches with similar experimental elbow and paw trajectories that were chosen from mouse behavior dataset. The mean synthesized excitations are shown in thick red for biceps long head, thick cyan for triceps long head, and thick blue for triceps medial head compared to the base lateral head activity. **B**. Violin plots including the entire dataset of 60 reaches. The mean synthesized muscle activity lies largely within 1 standard deviation of mean experimental muscle activity (red dots). On a reach-by-reach basis, the synthesized muscle activity lies largely within a z-score of 2 standard deviations (blue violin plots). Black box plots denote median z-deviation (white bar), 25 to 75^th^ percentile distributions (black box), and 10^th^ to 90^th^ percentile distributions (horizontal black line). **C**. A comparison of mean absolute error between time-shuffled physiological EMG data and synthetic excitation means to the real mean of the tracked data. Synthetic excitation means have lower MAE than the time-shuffled data in all three muscles recorded (two-sided t-test, biceps p = 3.1e-6, triceps long head p = 2.7e-6, triceps lateral head p =2.4e-3. P-values were Holm-Bonferroni corrected for multiple comparisons). Black box plots denote median (red horizonal bar), 25 to 75th percentile distributions (black box), and 10th to 90th percentile distributions (short horizontal black lines and stems).

**Figure 4.**
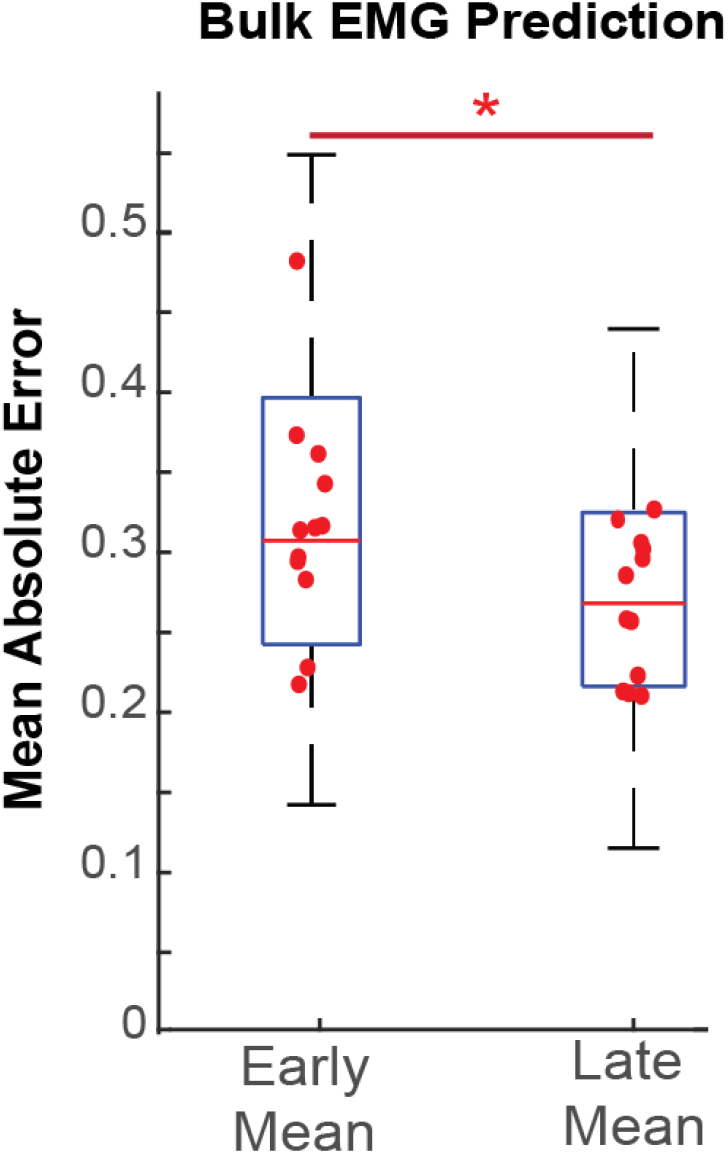
A comparison of the utility of the simulated muscle excitations in predicting EMG during early or late training sessions (i.e. the first three sessions after the first successful reaches being *early* and sessions later than four sessions after the initial successful reach being *late*.) Late reaches, on bulk, have a significantly lower mean absolute error than early sessions (two-sided t-test, p = 3.5e-6, early N = 40, late N = 60) for predicting trial-to-trial EMG activity. Black box plots denote median (red horizonal bar), 25 to 75th percentile distributions (black box), and 10th to 90th percentile distributions (short horizontal black lines and stems).

## RESULTS

### Kinematic Tracking

We tasked the physiological forelimb model to track recorded DeepLabCut-tracked kinematics using optimal control algorithms (see **Methods: Optimal control)**. The model was scaled and then optimized with direct collocation to track the paw and elbow across the ballistic epoch of the reach (Fig. 2 AB). Animals have natural variance in produced reaching movements and motor control, so we opted to group six sets of ten reaches by their kinematic similarity across time (Fig. 2C; see **Methods: Selection of reaches for simulation)**. We deliberately selected these sets of varying reaching kinematics to explore the ability of the model to predict varied motor behaviors. We were able to recreate limb kinematics with low error, with the majority of synthesized kinematics per timestep falling within 1 standard deviation of the empirical kinematic mean (Fig. 2D, blue violin plots; N = 60 reaches) across the x, y, and z dimensions of the paw and elbow trajectories.

### Model muscle activity patterning for reaching movements

We next tasked the model to synthesize muscle activity during reaching movements with optimal control in order to predict the muscle excitations that underlie observed kinematics. The predictions were not trained with ‘ground truth’ or empirical muscle activity. As shown in the examples in Fig. 3A, we observe that the mean synthesized muscle activity closely resembles empirical muscle activity over the duration of the reach for all three muscles. We measure the performance of the model via the mean absolute error (MAE) of normalized ground truth EMG signals from model signals. The mean muscle excitations produced by the model were typically within a single standard deviation of the experimental EMG activity (Fig. 3B, red dots in violin plots; between 50-57 of the 60 reaches, depending on the mice and assayed muscle).

On a reach-by-reach basis, the muscle excitations produced by the model across all time points were typically within two standard deviations of the experimental EMG activity, suggesting that, while still relatively accurate, individual reaches are more difficult to accurately predict. Paired reach-to-reach predictions were likely less accurate because mice were highly variable in their muscle patterning, even between reaches with the same kinematic profile (Supp. Fig 1). Some of the observed muscle patterning may be energetically inefficient, more consistent with early learning or motor exploration, which would not be predicted as accurately by optimal control approaches. In Fig. 3C, we show that the mean model EMG predictions outperform the shuffled experimental EMG data (i.e., having the same distribution as the ground truth EMG; two-sided t-test, biceps p = 3.1e-6, triceps long head p = 2.7e-6, triceps lateral head p =2.3e-3.; P-values were Holm-Bonferroni corrected for multiple comparisons, N = 60 shuffled trials and six synthesized means).

### Mouse motor learning and optimal control

The progression of reach kinematics and muscle patterning in mice learning a novel task is a relatively understudied phenomenon, especially given the paucity of experimental approaches to studying whole-limb muscle activity. We leveraged our model’s explanatory power to investigate the possibility that mice approach an optimal motor control solution during the progression of training by evaluating optimal control predictions during early and late sessions of training. In Fig. 3, we compared the optimal control predictions with reaches selected from expert mice (i.e., after at least 5 sessions of successful reaching). In Fig. 4, we compared how the optimal control predictions varied when the reaches were chosen in the early (i.e., in the first 3 sessions after the first successful reach to pellet) or late stages of learning. We found that mice tended to use muscle excitation patterns that converged more closely to those derived from optimal control in the later stages of learning. These results were significant when pooling the data across all recorded EMG channels but not on individual channels, likely because of our small sample size (early N = 4, late N = 6 for means comparisons, early N = 40, late N = 60 for trial-to-trial comparisons shown in Fig. 4. Comparison of trial-to-trial data was compared with a two-sided t-test with a p-value of 3.5e-6).

## DISCUSSION

Mouse models are widely used to study the neural control of movement, motor disorders, muscle physiology and develop novel brain-computer interfaces and neurotechnologies. Despite the widespread use of mice in the health sciences, the only available biomechanical model of the mouse forelimb is based largely on educated guesses, which could lead to inaccurate kinematics and muscle activity predictions. We used high-detail anatomical reconstruction from large scale light sheet microscopy scans to develop the first physiological biomechanical model of the mouse forelimb in terms of musculoskeletal geometry and muscle architecture (Mathewson et al., 2012). Traditional dissection methods were deemed impractical to determine the musculoskeletal geometry because of the small size of the mouse forelimb, especially in determining the attachment points of the tendons of the elbow. Other imaging techniques such as microCT would have likely also been adequate to produce sufficient soft tissue contrast for the reconstruction (Charles et al., 2016). We used this biomechanical model with optimal control to synthesize muscle coordination patterns that produce reaching movements in simulations that match experimental kinematics. Accurately predicting muscle activity is challenging because of the infinite possible coordination patterns consistent with the tracked experimental kinematics and the high physiological variance in the patterns observed in real mice (i.e., for very similar kinematics, mice often use very different muscle patterning strategies, some of which may be energetically costly, have high or low co-contraction, be robust to disturbances, etc.; Supplemental Figure 1). Our optimal control cost function only has terms to encourage low energy (via the sum of muscle activations squared proxy) and producing kinematics consistent with experimental kinematics. Therefore, we would not expect the optimal control predictions to closely match the experimental muscle activity on a reach-by-reach basis because of the high variability in the experimental muscle patterning data. Nevertheless, we found that the mean optimal control muscle activity predictions have strong resemblance with the mean empirical muscle activity (Fig. 3A). These results held for all three recorded muscles with EMG (biceps, triceps long head and triceps lateral head). To the best of our knowledge, this is the first work in any species, including humans, showing resemblance between synthesized and experimental muscle activity for three-dimensional reaching movements with a biomechanical model.

Neuroscience experiments are sometimes limited in scope by the difficulty of simultaneous recording of behavior, neurological signals, and, in some cases, muscle activity. Multisite muscle recordings are limited to accessible sites, and this limitation is exacerbated in mice, where access to and implantation of many muscles is often infeasible. This model is meant to supplement experiments where knowledge of muscle activity patterning could bring insight about the nature of neural activity patterning. Scientists with behavioral data can extract an estimate of whole-forelimb muscle activity from the model given a set of kinematics over time. Tracking of mouse kinematics has become broadly accessible through the advent of pose-based tracking software like DeepLabCut, which was used in the present study to monitor limb position during reaching behaviors (Mathis et al., 2018). The conjunction of tracking and synthesis of whole-forelimb muscle activations promises to expand research into behavioral control significantly.

There are several extensions to our biomechanical model and computational tools possible for future work. Our computational tools assume that no EMG is available during the experiments. If EMG is collected during the experiment, the optimal control problem can be solved to predict the muscle activity of muscles without EMG recordings while matching experimental EMG and kinematics data in tracking simulations (Dembia et al., 2021). It is also possible to change the cost function in the optimal control problem and produce predictive simulations that do not require any experimental data, including kinematics, by specifying goals constrained only to endpoint or end-state of the limb. The optimal control problem could then predict the reaching kinematics and muscle activity when there is a change in the task (e.g., a new pellet location) or limb biomechanics (e.g., a weight placed on the forelimb). This model might also be used to assay disease states to expand on known motor symptoms, such as enforced variability in endpoint accuracy as seen in cerebellar disease (Bonnefoi-Kyriacou et al., 2018), forced variation in muscle coordination patterns such as those seen after stroke (Cheung et al., 2012) or replications of impaired kinematics in Parkinson’s disease (Vissani et al., 2021).

While the model evaluation in this study focuses on reaching movements, which are commonly used to assess motor control across species, the insights gained extend beyond this specific task. Reaching movements provide a well-studied framework for understanding motor coordination, but motor systems are capable of generating numerous possible muscle coordination patterns for any given movement (Harris & Wolpert 1998). Optimal control chooses the muscle excitation pattern that achieves task constraints, while minimizing a proxy for effort or energy (e.g., the sum of muscle activations squared) and possibly other terms (Al Borno et al., 2020). We apply optimal control to predict an energetically efficient muscle activity pattern that achieves the reaching kinematics task. This study is focused on the ballistic phase of reaching movements, and we did not model the grasping phase during the reach as we would have needed to include the muscles that control the wrist and fingers in the model and simulate interaction with the pellet. One additional discrepancy between our simulation and the empirical reach is that mice, before starting their reach, rested their paws on a bar, which we did not simulate (and may impact the predicted muscle activity at the start of the ballistic phase).

Our simulations were evaluated with head-fixed mouse reaching. Using the biomechanical model in free-reaching mice may be less accurate because it has more significant scapular movements, which we assume to stay fixed in our model. In future work, researchers could either add a degree-of-freedom and a joint motor to allow translational movement of the scapula or incorporate the muscles that control the scapula as a free body in the biomechanical model. The optimal control solutions produce open-loop muscle coordination patterns that are not responsive to noise or changing task or environmental constraints. It is, however, possible to develop closed-loop controllers to track the optimized trajectory or to develop feedback controllers with reinforcement learning or introduce stochastic noise representing imprecise neural controls (e.g., Van Wouwe et al., 2022). Finally, we generalized our muscle parameterization to past work and observations from other small mammals (Powell et al. 1984), but the model would certainly benefit from a thorough investigation of the murine forelimb’s muscles through dissection and testing, which was beyond the scope of our study.

We make our computational tools freely available as open-source. We have written custom code to convert our OpenSim model into the MuJoCo physics simulation environment (Todorov et al., 2012). Although the model is available in MuJoCo, the computational tools for optimal control are based in OpenSim; therefore, MuJoCo users will need to develop their own code to produce the simulations with the model. While the model evaluation in this study focuses on reaching movements, our tools can be used to predict muscle activity in other forelimb movements. Users of our computational tools should note that the optimal control predictions are expected to more closely resemble empirical muscle activity on a mean-basis rather than on a trial-by-trial basis and carry the assumption of closely matched kinematics. Furthermore, the predictions are expected to improve when mice have learned to perform the task well as opposed to when mice are still in the early stages of learning. Nevertheless, the model predictions in the early stages of learning are still within one standard deviation of empirical results and represent a significant improvement over randomized guesses from the naturalistic EMG distribution. An exciting use case for our biomechanical model is to control it with artificial neural networks and relate the activity in these networks with empirical neural activity from system neuroscience laboratories (Aldarondo et al., 2024). Combining our computational tools and experimental data could lay the foundations for future studies elucidating the principles that drive the control of movement.

## ACKNOWLEDGEMENTS

JIG designed the study, wrote the manuscript, assembled the model, and performed analysis. AGC performed imaging and prepared mice under the supervision of DH.. SKC designed the behavioral experiment, performed the headplate and EMG surgeries, trained the mice, recorded electromyography, and provided datasets under the supervision of AP. JH helped design and conceptualize the study. MAB designed the study, wrote the manuscript, provided tools for analysis and supervised the study. We thank InWorks at CU Denver for imaging and segmentation. The behavioral study included in this work was supported by NIH grants (along with Simons Consortium): U24NS126936 and R01NS109237. JIG was supported by NLM grants 2T15LM009451-16 and 5T15LM009451-15.

---

**Supplemental Figure 1.**
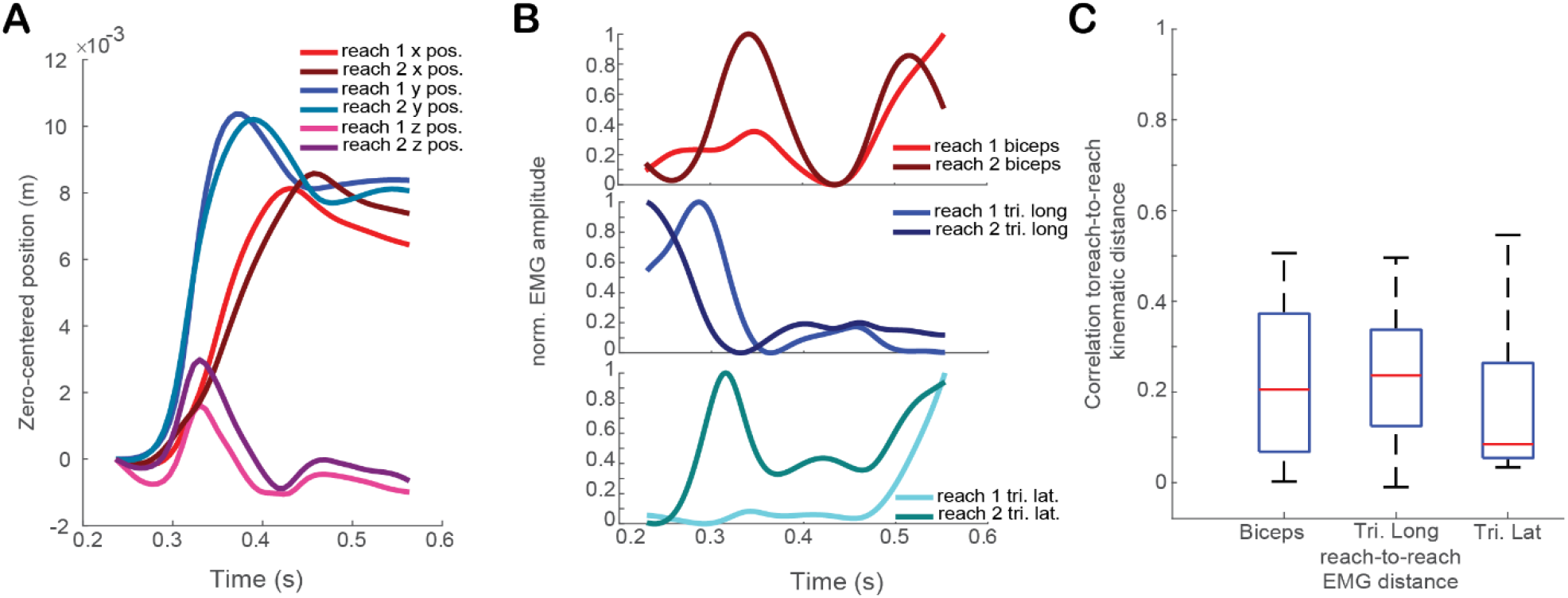
Variability in electromyography between similar reaches. **A**. Example of two reaches that have similar trajectories, with x from teaches one and two shown in red, y in blue, and z in magenta. Reaches have been aligned to their initial position. **B**. Corresponding EMG signals from the reaches selected in A, with biceps activity for reaches one and two shown on the top panel in red, triceps long head in blue in the second panel, and triceps lateral head shown in teal in the third panel. **C**. Correlation between the Euclidean distance between reach trajectories to the distance between normalized EMG traces, showing a low to moderate effect on EMG similarity for alike reaches.

