## Supplemental Table 1 for "A novel biomechanical model of the mouse forelimb predicts muscle activity in optimal control simulations of reaching movements"

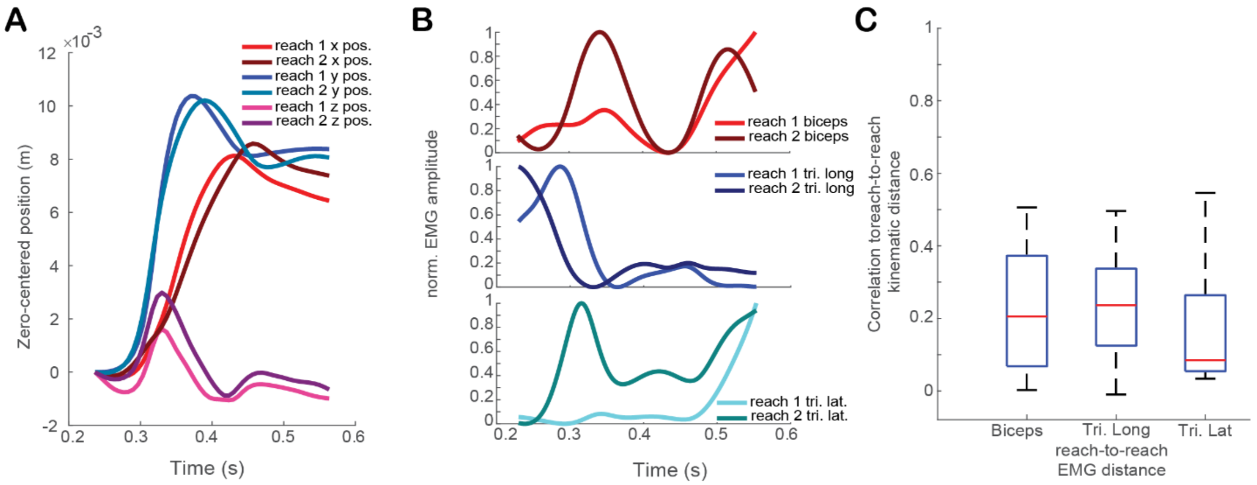


**Supplemental Figure 1. Variability in electromyography between similar reaches. A.** Example of two reaches that have similar trajectories, with x from teaches one and two shown in red, y in blue, and z in magenta. Reaches have been aligned to their initial position. **B.** Corresponding EMG signals from the reaches selected in A, with biceps activity for reaches one and two shown on the top panel in red, triceps long head in blue in the second panel, and triceps lateral head shown in teal in the third panel. **C.** Correlation between the Euclidean distance between reach trajectories to the distance between normalized EMG traces, showing a low to moderate effect on EMG similarity for alike reaches.

Supplemental Table 1. Simulation Computer Specifications

| | |
|---|---|
| Processor | 12th Gen Intel(R) Core(TM) i9-12900K, 3200 Mhz, 16 Core(s), 24 Logical Processor(s) |
| BIOS Version/Date | Dell Inc. 2.13.0, 3/8/2024 |
| System Model | Dell Precision 3660 |
| OS Name | Microsoft Windows 11 Pro |
| Installed Physical Memory (RAM) | 128 GB |
